## Supplementary Data for "Arrayed CRISPRi and Quantitative Imaging Describe the Morphotypic Landscape of Essential Mycobacterial Genes"

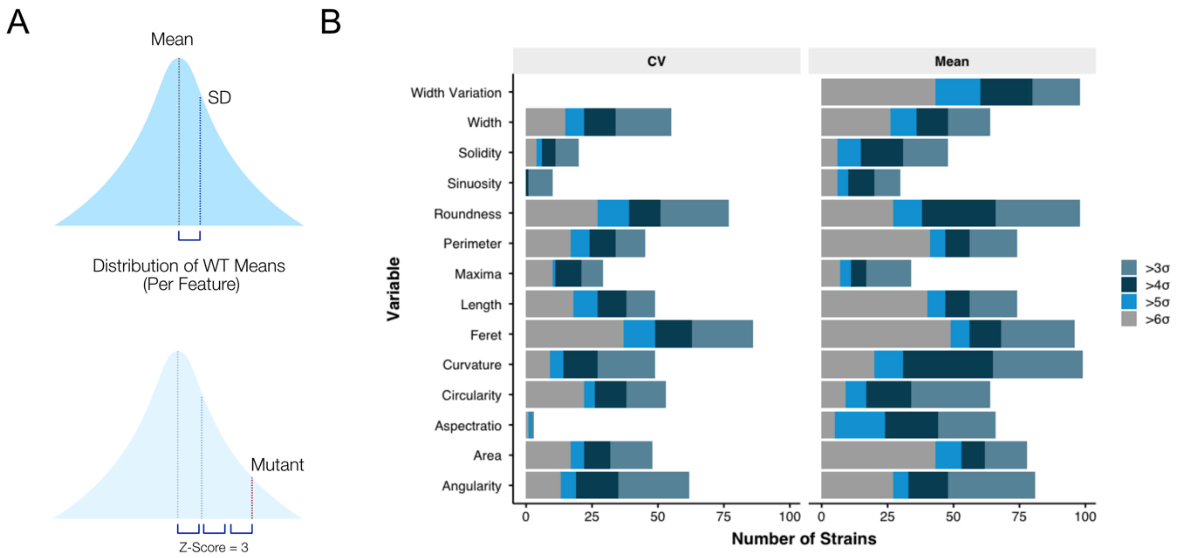


***Supplementary Figure 1.*** *The morphological impact of essential gene silencing. (A) Means and Coefficients of Variation (CV) were measured for each morphological feature, for each gene knockdown, followed by a Z-score transformation. (B) From the arrayed library, 206 (78%) mutants had at least one morphological feature with a Z-score >3, or <-3.*

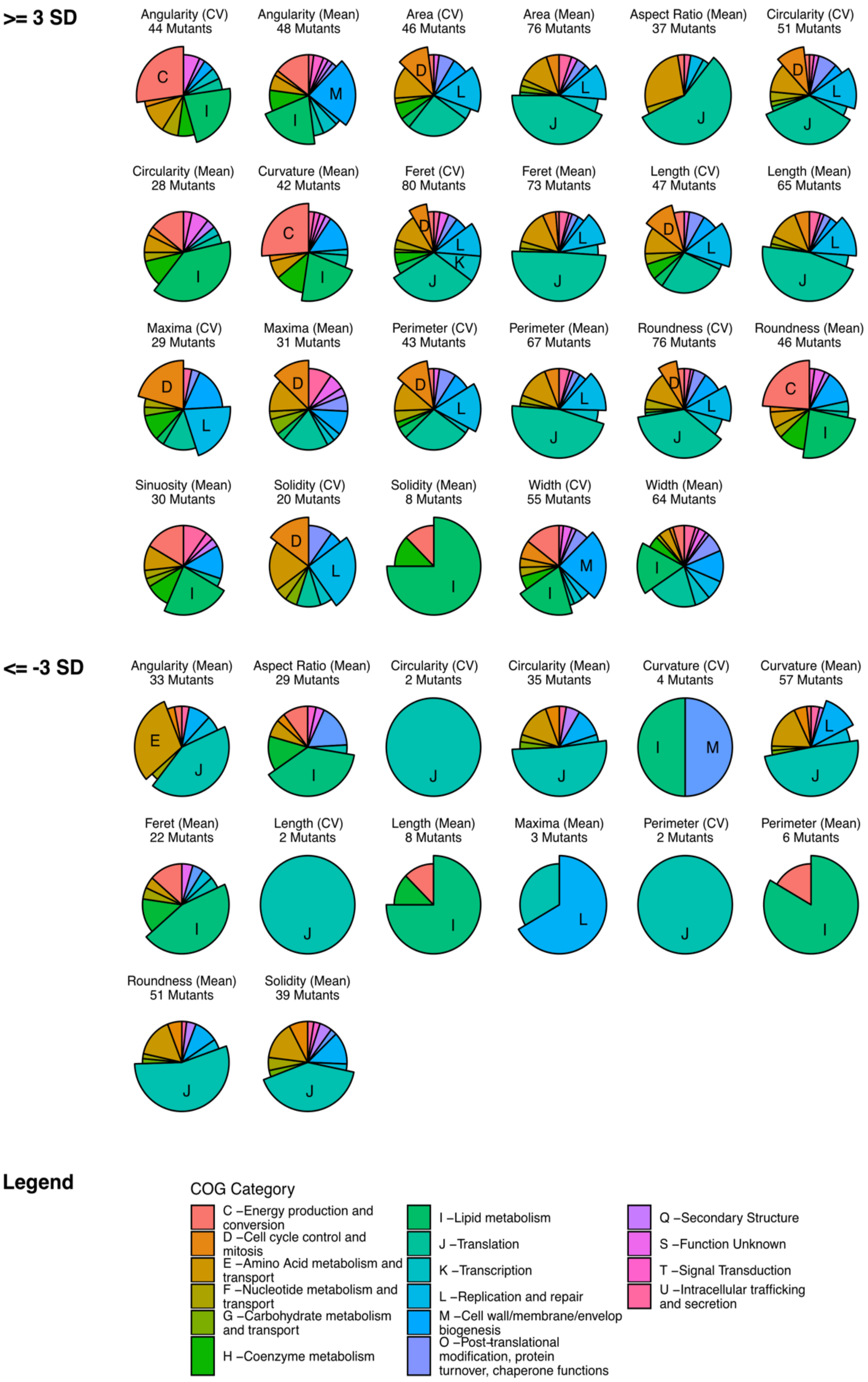


***Supplementary Figure 2.*** *COG enrichment identifies associations between genetic function and morphological changes. Each gene in the library was functionally annotated to a Cluster of Orthologous Groups (COG) category. For each morphological feature, mutants were selected with Z-scores >3, or<-3, and tested for statistical enrichment of COGs, with adjustment for multiple testing. Only features with at least one statistically enriched COG (adjusted p value ≤ 0.05) are shown and labelled.*

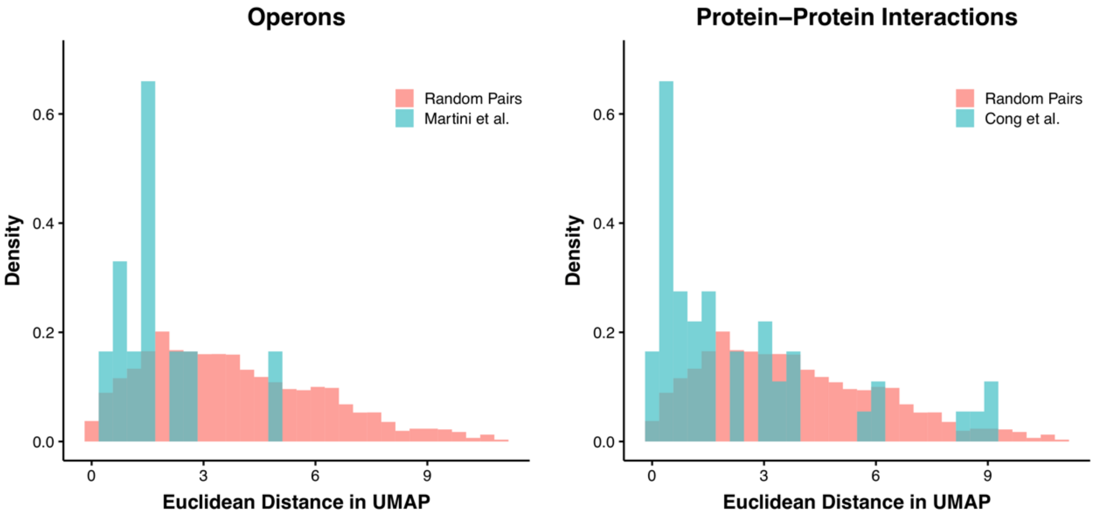


***Supplementary Figure 3.*** *Distances in UMAP space can reflect relationships of biological relevance. We obtained data on high-confidence operon identification from a published report (Martini, Zhou et al. 2019) and compared the Euclidean distances of genes in identified operons to random pairs of genes. A similar approach was used for published protein-protein interaction predictions (Cong, Anishchenko et al. 2019).*

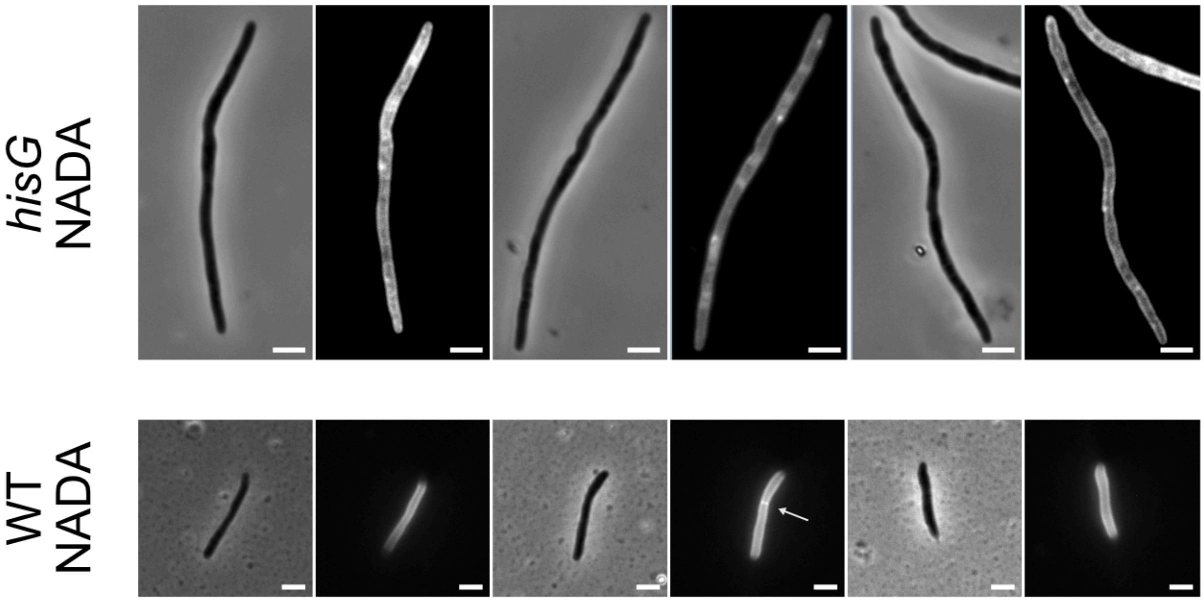


***Supplementary Figure 4.*** *Absence of septum formation on histidine depletion. Following CRISPRi-mediated silencing of hisG, cells were stained with the D-alanine analogue, NADA, for visualization of peptidoglycan (Botella, Yang et al. 2017). While staining was uneven, no clear septa were detected in any cells imaged (n = 35).*

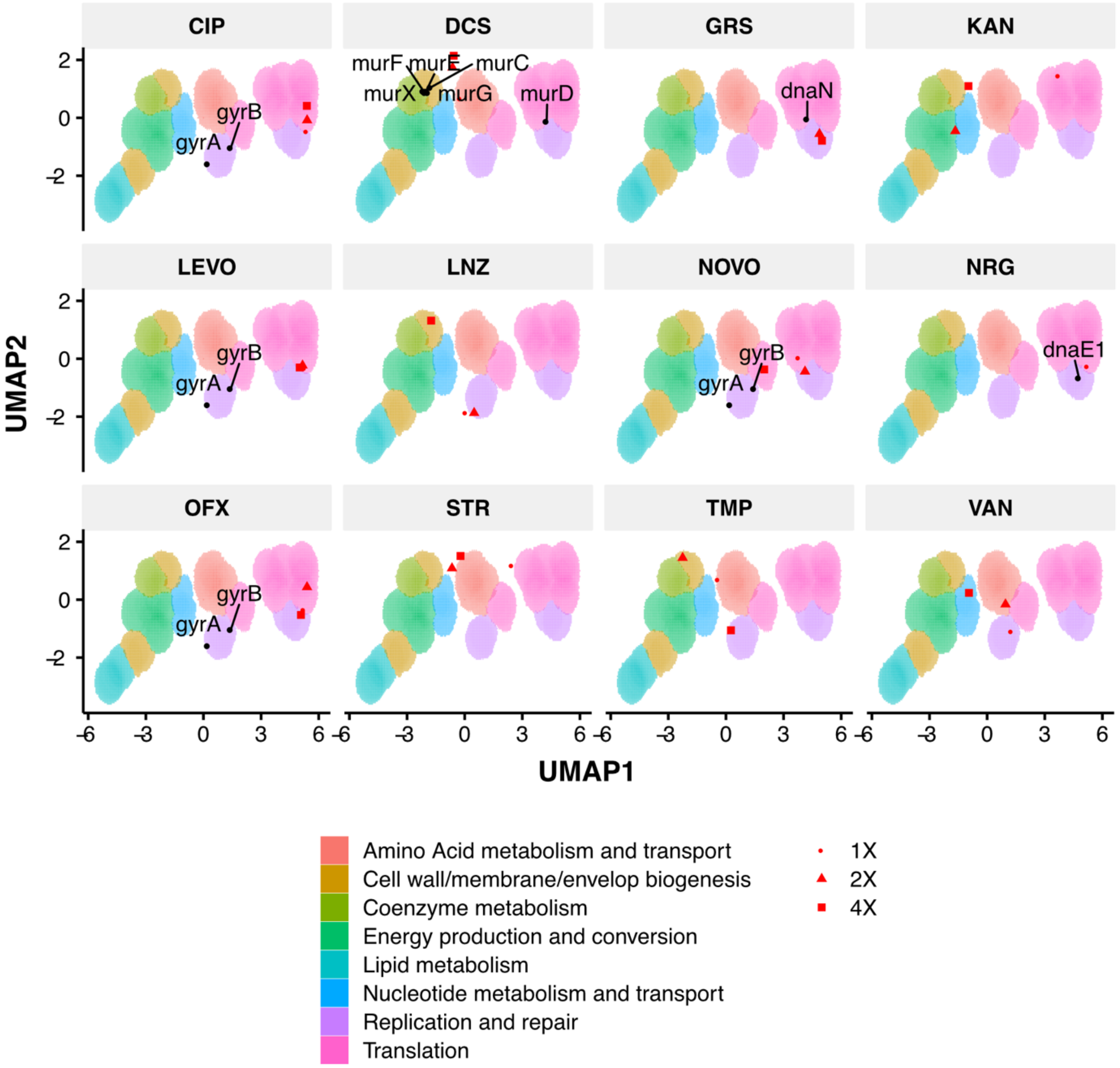


***Supplementary Figure 5.*** *Phenoprinting can inform antimicrobial MOA. Cells were exposed to varying supra-MIC concentrations (1X, 2X, 4X MIC) of the selected antimycobacterial compounds for 18 hours, imaged, and analyzed using the same pipeline developed for CRISPRi imaging. The resulting profiles were visualized in CRISPRi-generated UMAP space. CIP, ciprofloxacin; DCS, D-cycloserine; GRS, griselimycin; KAN, kanamycin; LEVO, levofloxacin; LNZ, linezolid; NOVO, novobiocin; NRG, nargenicin; OFX, ofloxacin; STR, streptomycin; TMP, trimethoprim; VAN, vancomycin.*

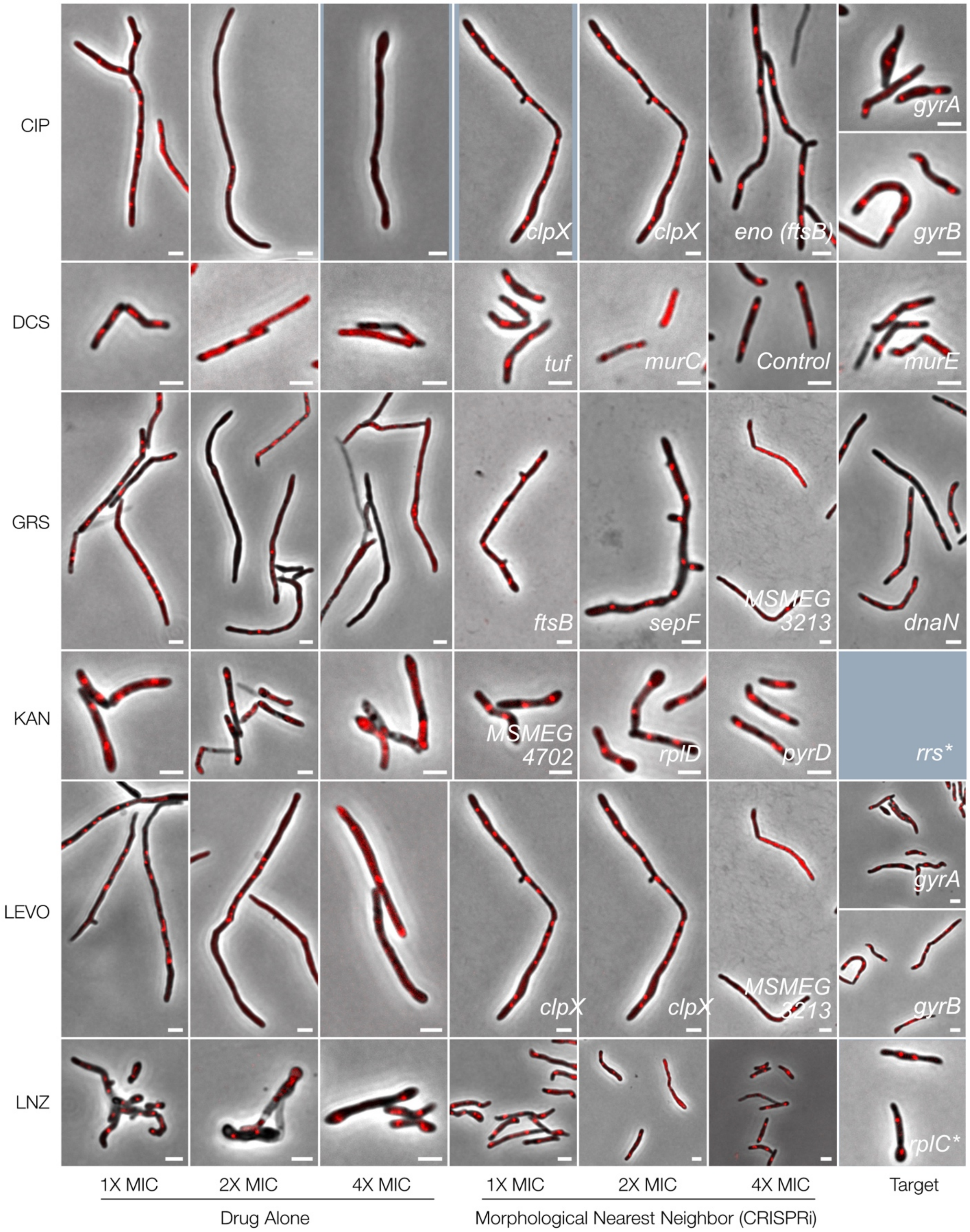
CIP
DCS
GRS
KAN
LEVO
LNZ
clpX
clpX
eno (ftsB)
gyrA
gyrB
tuf
murC
Control
murE
ftsB
sepF
MSMEG 3213
dnaN
MSMEG 4702
rplD
pyrD
rrs*
clpX
clpX
MSMEG 3213
gyrA
gyrB
rplC*
1X MIC
2X MIC
4X MIC
1X MIC
2X MIC
4X MIC
Target
Drug Alone
Morphological Nearest Neighbor (CRISPRi)

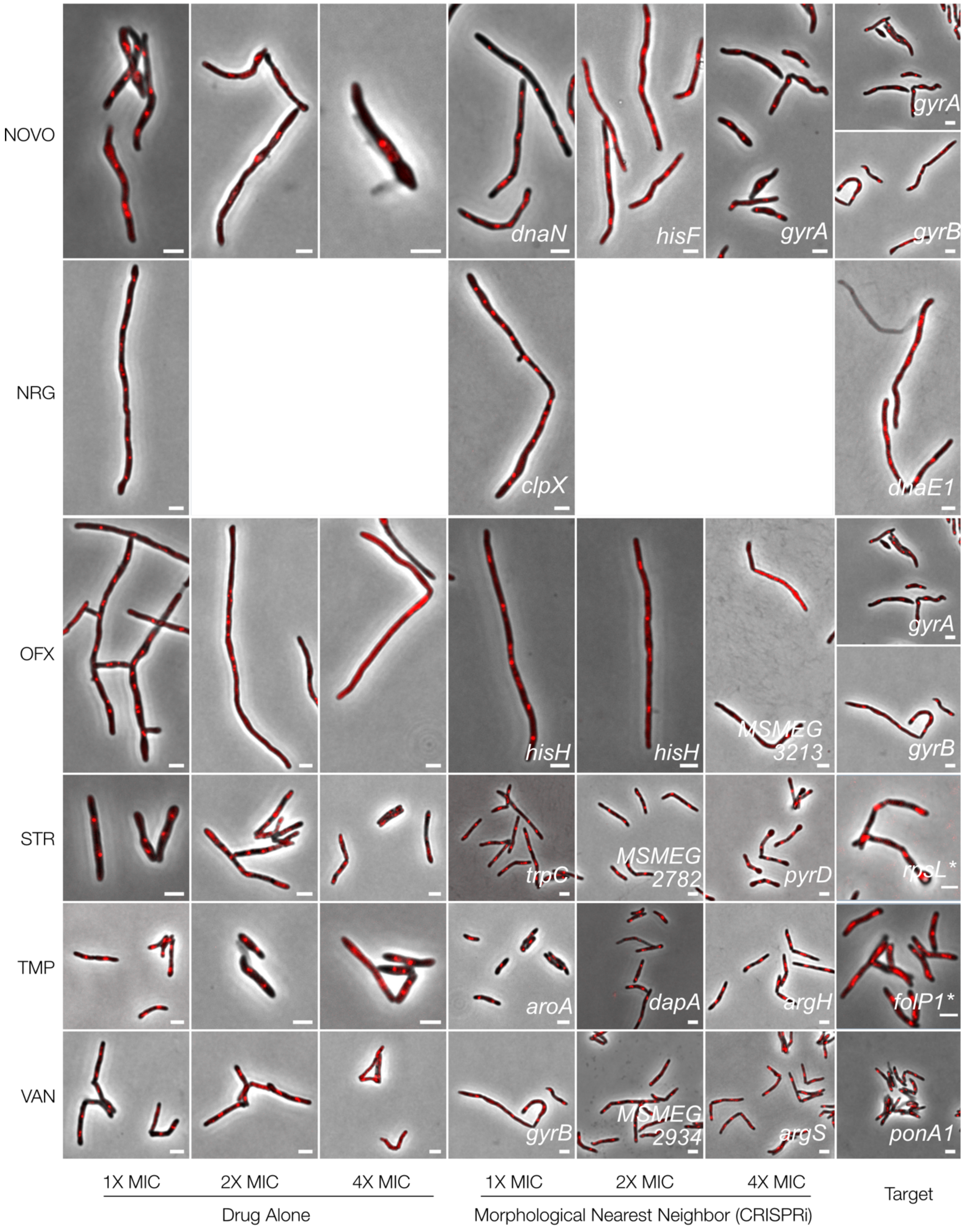
NOVO
NRG
OFX
STR
TMP
VAN
dnaN
hisF
gyrA
gyrA
gyrB
clpX
dnaE1
hisH
hisH
MSMEG 3213
gyrA
gyrB
trpC
MSMEG 2782
pyrD
rpsL*
aroA
dapA
argH
folP1*
gyrB
MSMEG 2934
argS
ponA1
1X MIC
2X MIC
4X MIC
Drug Alone
1X MIC
2X MIC
4X MIC
Morphological Nearest Neighbor (CRISPRi)
Target

***Supplementary Figure 6.*** *Phenoprinting can inform antimicrobial MOA.* ***.*** *Cells were exposed to varying supra-MIC concentrations (1X, 2X, 4X MIC) of the selected antimycobacterial compounds for 18 hours, imaged, and analyzed using the same pipeline developed for CRISPRi imaging. The resulting profiles were visualized in CRISPRi-generated UMAP space. For each compound, representative cells, known targets and the morphological nearest neighbor are presented. CIP, ciprofloxacin, DCS, D-cycloserine; GRS, griselimycin; KAN, kanamycin, LEVO, levofloxacin; LNZ, linezolid. NOVO, novobiocin; NRG, nargenicin; OFX, ofloxacin; STR, streptomycin; TMP, trimethoprim; VAN, vancomycin.*

**The major spontaneous resistance gene, if available. For kanamycin, rrs is not present in the library. Limited compound availability restricted nargenicin assays to 1X MIC.*

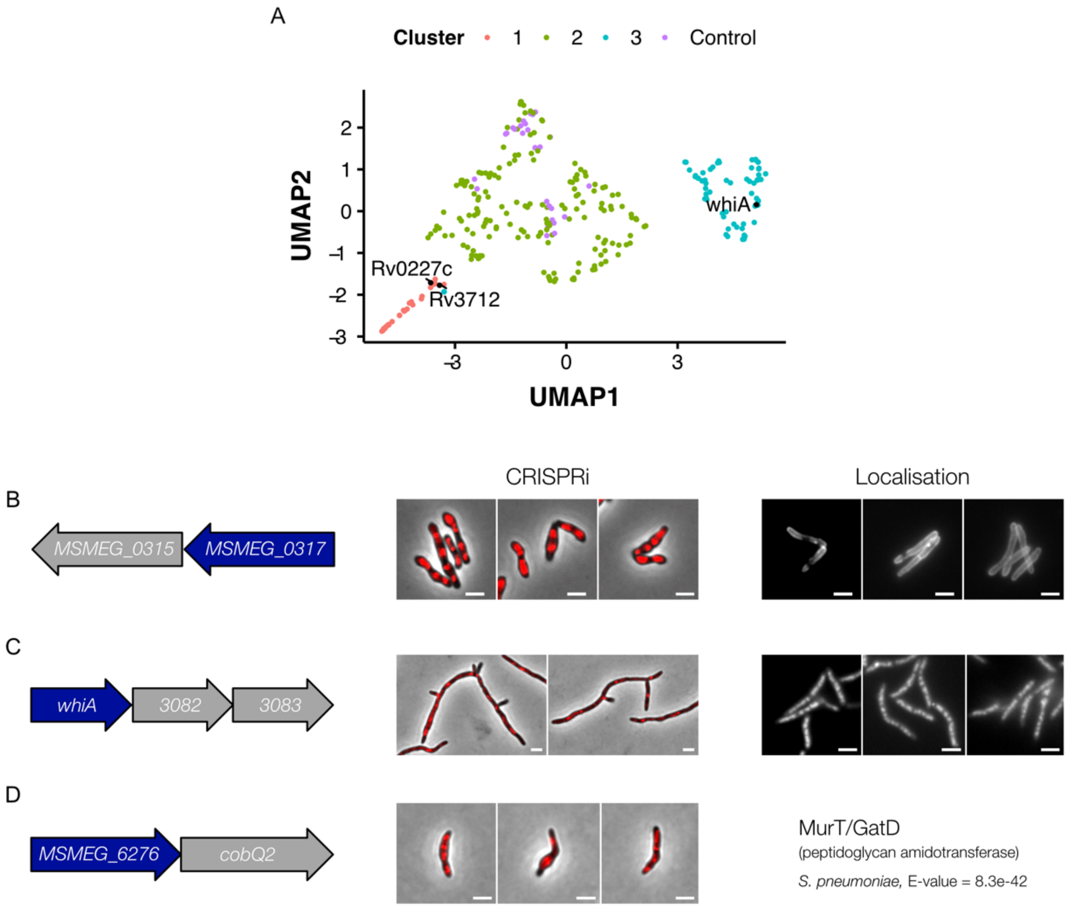


***Supplementary Figure 7****. Morphological profiling informs gene function. (****A****) Genes with putative function were visualized in UMAP space. MSMEG_0317 (Cashmore, Klatt et al. 2017) and MSMEG_6276 (murT) were found in the cell wall cluster, supporting their suggested roles as components of cell wall synthesis. The transcription factor, WhiA (Rustad, Minch et al. 2014), was found in the DNA-Cell Division-Translation Cluster, suggesting a role in cell-cycle regulation. (****B****) MSMEG_0317 produces a distinct phenotype on knockdown. Dendra-tagged MSMEG_0317 localizes to the cell wall and cell septa. (****C****) whiA knockdown produces elongated, branching cells. Dendra-tagged WhiA is localized to the nucleoid. (****D****) MSMEG_6276, a gene with strong homology, according to HHpred (Hildebrand, Remmert et al. 2009), to murT/gatD produces bulging and lysis on silencing.*

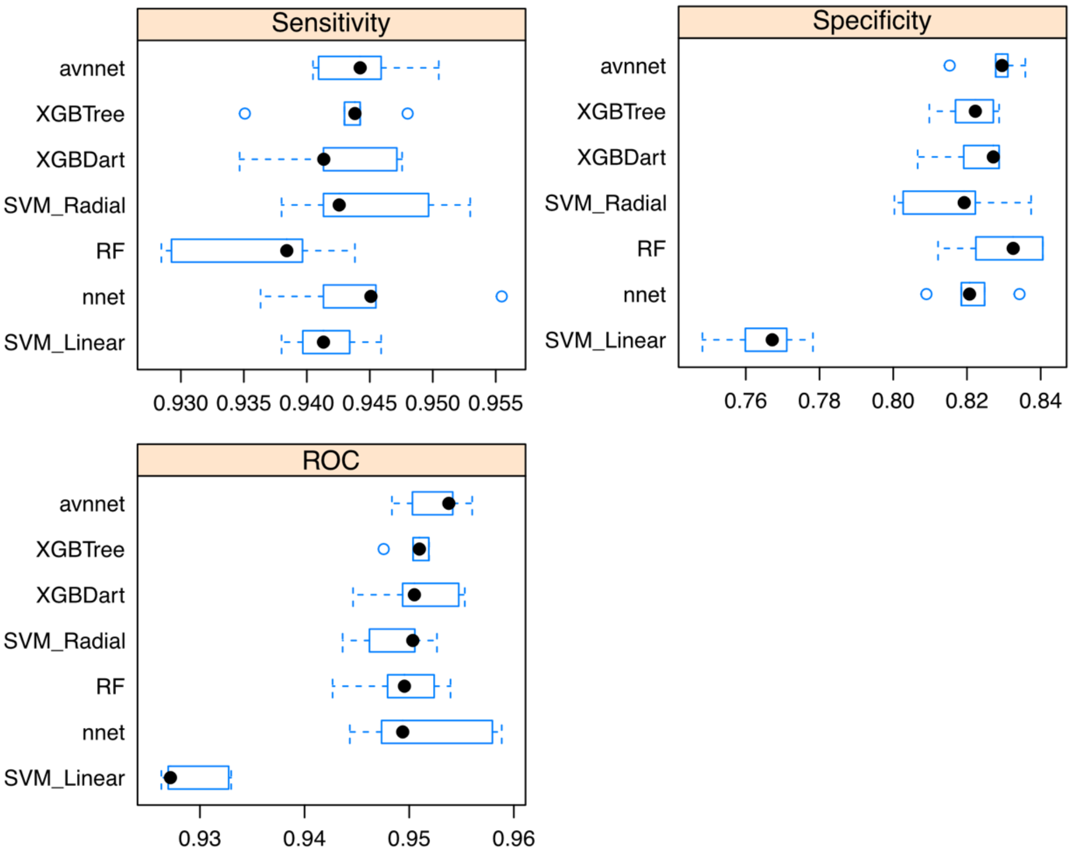


***Supplementary Figure 8.*** *Comparisons of cell classifier models. Samples of the imaged library were analyzed with MicrobeJ and manually classified. A variety of models was trained using Caret, with 5-fold cross validation, and tested on a reserved sample of the classified images. An averaged neural network was selected as the best performing model based on Receiver-Operating Characteristic (ROC) area-under-the-curve.*

### SUPPLEMENTARY METHODS

#### Sequencing PCR Reaction

Sequencing Primers (5’ to 3’):
Forward: CTGGTCCACCTACAACAAAG
Reverse: CCCTGATTCTGTGGATAACC

| STEP | TEMP | TIME |
|---|---|---|
| Initial Denaturation | 94°C | 30 seconds |
| 30 Cycles | 94°C<br>49°C<br>68°C | 15-30 seconds<br>15-60 seconds<br>1 minute/kb |
| Final Extension | 68°C | 5 minutes |
| Hold | 4-10°C | |

#### Golden Gate PCR Reaction

Golden Gate Primers (5’ to 3’)::
Forward: ACTTCGGCTCTTCG*GGA*TCTGACCAGGGAAAATAGC
Reverse: ACTTCGGCTCTTCGCTGAAAATAAAAAAGGGGACCTCTAG

| STEP | TEMP | TIME |
|---|---|---|
| Initial Denaturation | 98°C | 2 minutes |
| 30 Cycles | 98°C<br>50°C<br>72°C | 10 seconds<br>30 seconds<br>30 seconds |
| Final Extension | 72°C | 3 minutes |
| Hold | 4 | |
